# ROLE OF WORKING MEMORY IN INTERLIMB GENERALIZATION OF NEWLY LEARNED SKILLS

**DOI:** 10.1101/2024.08.17.608389

**Authors:** Goldy Yadav, Rahul Pal, Shraddha Matkar, Neeraj Kumar

**Author notes:** These authors contributed equally to this work (Goldy Yadav, Rahul Pal).

## Abstract

Newly acquired skill memory can generalize/transfer to the untrained arm. Such interlimb generalization of a learned skill has been shown to be symmetric in nature and is thought to be mediated by cognitive processes that emerge during skill learning. However, it is unknown whether engaging in other cognitively demanding tasks following skill acquisition can influence skill generalization. Our research goal was to uncover how a secondary task, involving working memory, interacts with a newly formed skill memory and influences subsequent interlimb generalization. To test this idea, we conducted a set of three experiments by recruiting right-handed young healthy individuals (N=92) who learned a novel motor skill (long or short training on a skilled reaching task) followed by performing a working memory or control task with the right arm. Finally, all individuals were tested for immediate or delayed (after 24 hours) interlimb skill generalization to the untrained left arm. We found significant immediate as well as delayed generalization in individuals who received long training on the motor skill task, irrespective of whether they performed working memory or control task. On the other hand, performing the working memory but not control task following short skill training impaired generalization when the untrained arm was tested 24 hours later. These findings indicate that short training reflecting early stages of skill learning and the subsequent skill memory stabilization are dependent on working memory such that the underlying neural interactions mediating these processes can have implications for skill generalization.

**NEW AND NOTEWORTHY:** Skill learning is deemed successful if it can generalize to untrained conditions. Despite the potential clinical relevance of interlimb skill generalization for motor rehabilitation, such as in case of unilateral stroke, the underlying mechanisms that can influence such generalization are not fully understood. Specifically, whether interlimb skill memory generalization can be impacted by another memory system is unknown. Here, we show that engaging in a working memory task following short skill training is detrimental for interlimb skill generalization, and such interference caused by working memory is not evident when the skill training is longer. These behavioral findings bridge the link between traditionally distinct memory systems, i.e., procedural skill memory and working memory.

## INTRODUCTION

Motor learning is a unique aspect of human behavior and a critical research area in psychology and neuroscience. Typically, motor learning abilities are characterized by acquisition of novel skilled movements (motor skills) and refinement of the existing ones (motor adaptation). Motor skill learning leads to reduced movement errors, improved movement speed and accuracy, and eventually results into stable memory formation of the newly learned movements (Robertson et al., 2009; Reis et al., 2009; Shmuelof et al., 2012; Telgen et al., 2014; Yadav and Duque, 2023). On the other hand, learning during motor adaptation requires correcting for movement errors resulting from an external perturbation (Shadmehr and Mussa-Ivaldi, 1994; Thoroughman and Shadmehr, 2000, Taylor, Krakauer and Ivry, 2014). Both motor skill learning as well as motor adaptation are deemed successful if such abilities can generalize or transfer to untrained contexts (Poggio and Bizzi, 2004; Krakauer et al., 2006). Understanding how learning generalizes to unpracticed conditions can provide deeper insights into the underlying human motor learning mechanisms.

One form of generalization that specifically interests us, given its relevance for motor rehabilitation, is interlimb generalization in which learning acquired using one effector such as the arm is transferred to the other untrained arm. Studies have shown that such interlimb generalization is possible, but it is variable and influenced by task conditions (Joiner et al., 2013) and individual movement characteristics (Renault et al., 2020). For example, many studies have shown that generalization of motor adaptation is asymmetric and unidirectional, i.e., from non-dominant arm to the dominant arm (Criscimagna-Hemminger et al., 2003; Wang and Sainburg, 2003, 2004; Bao et al., 2017; Kumar et al., 2018; Kumar et al., 2020). Moreover, factors such as workspace location (Wang and Sainburg, 2006), handedness (Inui, 2005; Chase and Seidler, 2008; Lefumat et al., 2015), movement speed (Lefumat et al., 2015) and the amount of training (Kumar et al., 2020) can influence such interlimb generalization of motor adaptation. In contrast, our previous work shows that generalization of newly learned motor skills is symmetric, bidirectional (i.e., from non-dominant to dominant arm and vice versa) and driven via transfer of explicit/cognitive strategies developed during training (Yadav and Mutha, 2020). However, factors that can influence such generalization of a newly acquired motor skill memory to the untrained arm (specially the non-dominant arm) are poorly understood. Intrigued by this, we asked a simple question-can generalization of a newly acquired skill memory be impacted by performing a post-skill training secondary task that is cognitively demanding?

Skill motor behavior is dependent on formation and stabilization of motor memories. However, newly formed memories are unstable, and liable to disruption and interference (Robertson et al., 2004; Cohen and Robertson, 2011; Robertson, 2009; 2012). Studies exploring the fate of a memory post-training suggest that instability of the memory provides a time window for interaction that may allow interference and impact information generalization (Seidler, 2004; Mosha and Robertson, 2016; Robertson, 2018). On the other hand, memory stability can be enhanced by increasing training duration as experts who have mastered a skill train repeatedly to stabilize that skill and make it less vulnerable to disruption by interference. Based on such an interplay between memory stability and interference, we probed whether skill generalization of a newly acquired motor memory after a long versus short training is influenced by engaging in a secondary memory task (such as working memory). To test this idea, we conducted a series of experiments in which young healthy individuals trained on a novel motor skill task (variable task structure requiring fast and accurate movement to random targets) using their dominant right arm. Following training, one set of individuals performed a visuo-spatial working memory task known to be mediated by neural substrate such as dorsolateral prefrontal cortex (DLPFC) which is interestingly also involved in memory formation following variable skill training (Sala et al., 2003; Klauer and Zhao 2004; Fregni et. al., 2005; Sun et al., 2007; Kantak et al., 2010). The other set of individuals performed a control task with no working memory requirement with their right arm. Finally, we tested the untrained left arm of all individuals for immediate or delayed (24 hours later) interlimb skill generalization to help us uncover interactions between newly formed skill memory resulting from long versus short training and a cognitively engaging secondary task of working memory. Our results showed significant immediate and delayed generalization after long skill training regardless of the secondary tasks, but performing the working memory task after short training impaired generalization when untrained arm was tested 24 hours later.

## METHODS

### Participants

A total of 92 young healthy individuals with normal or corrected-to-normal vision (age range: 18–38 years, mean±SD: 22.663 ± 3.61 years, 38 Females) successfully completed the study. All participants gave written informed consent. This study was approved by the Institute Ethics Committee of the Indian Institute of Technology Hyderabad. All participants were right-handed (Edinburgh Handedness Inventory; Oldfield, 1971) and were naive to the experimental procedure (apparatus, paradigm, and purpose of the study). No individual reported any history of neurological, medical, or psychiatric abnormality before taking part in the study. All participants were given monetary compensation for taking part in this study.

### Study Apparatus

The experimental setup consisted of a digitizing tablet (GTCO CalComp) placed on a frame over which a computer screen (1920 × 1080 pixels) was mounted horizontally with a reflective mirror placed in-between (Figure 1a). Participants sat on an adjustable chair in front of this setup and the height of the chair was adjusted for every participant so that they could clearly see the stimuli reflecting on the mirror. Direct vision of the hand was blocked and a yellow dot (2 mm) indicating location of the handheld stylus was provided to guide participant’s movement during the experiment. The hand position was recorded at 120 Hz sampling rate on the digitizing tablet. The tasks were designed using Psychtoolbox on MATLAB (Brainard, D. H., & Vision, S., 1997).

**Figure 1:**
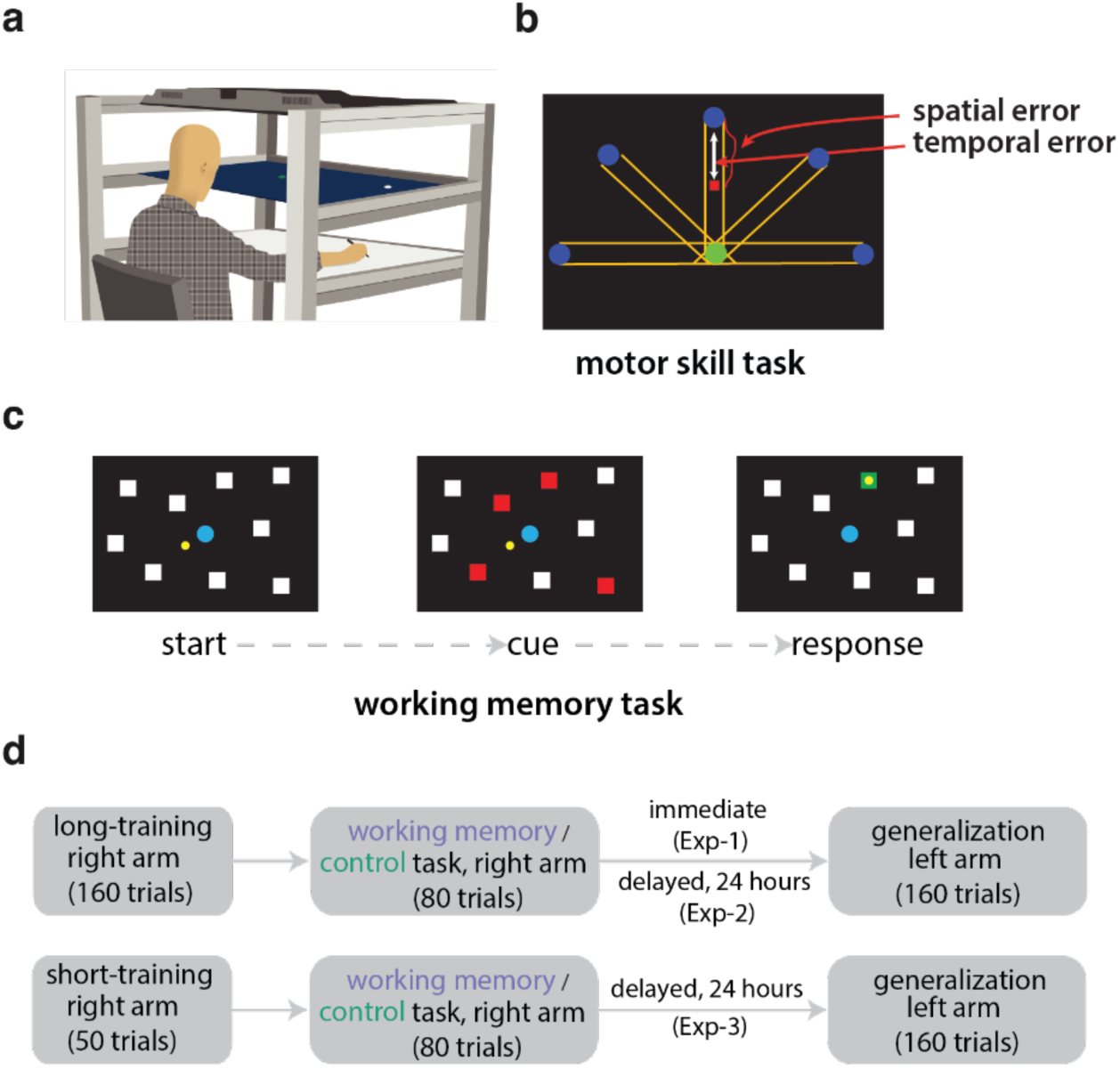
a) Setup with TV screen (top), reflective screen (middle) and digitizing tablet (bottom) on which individuals made planar movements and performed all tasks. b) Motor skill task with a representative movement from start (*green*) circle to one of the five target (*blue*) circles at a distance of 150 mm, connected by *yellow* lines. Note that only one target appears on each trial. Distance between the *red* square (indicating hand position at 650 ms of the executed movement) and target circle is considered *temporal error*. *Red* line indicating hand movement outside the *yellow* path is considered *spatial error*. c) Working memory task indicating start (presentation of nine *white* squares with *blue* start circle and *yellow* dot indicating hand position), cue (in this representation four out of nine squares changed from *white* to *red*. Note that the color change was sequential i.e., *white* squares flashed in *red* for 750 ms which was to be recalled later testing the working memory. However, for the control task the red squares were present on the screen for the entire trial duration), and the response screen (during which participants moved to touch the squares that turned *red*). d) Experimental design depicting Exp-1 (Long-Immediate) with long-training, secondary task (working memory, N=16 /control, N=16) with right hand, and testing untrained left arm for immediate interlimb generalization. Same for Exp-2 (Long-Delayed, N=30) except that generalization was delayed by 24 hours (working memory, N=15 /control, N=15). Finally, Exp-3 (Short-Delayed, N=30) contained short-training, secondary tasks and generalization at 24 hours (working memory, N=15 /control, N=15).

### Task Procedure

#### Motor Skill Task

In the beginning of the experiment all subjects had a brief familiarization session about the setup and the task. For the motor skill task (Figure 1b), participants were asked to perform fast and accurate planar reaching movement between a start circle (green circle, 10 mm diameter) and a target circle (blue circle, 10 mm diameter) with the right hand holding the stylus on the digitizing tablet. Each trial started with the participant moving his/her right hand to the start circle. After 500 ms at this start circle, an audio cue (beep) was presented, and one of five target circles appeared (pseudorandomized) on the screen. These two circles were 150 mm apart and connected with parallel yellow lines/path. Participants were instructed to move to the target circle as soon as they heard the audio cue while trying to remain within the yellow path. The yellow dot indicating the hand position was visible throughout the movement. Thus, to make fast and accurate skilled movements on this task participants were explicitly told to finish the movement as fast as possible (i.e., within 650 ms based on our pilot data; indicating temporal accuracy) and reach the target circle while moving within the yellow path (indicating spatial accuracy). Trial duration was 2 seconds and upon completion, movement performance feedback was provided through visual cues and a numerical score-visual cues comprised of a red line indicating part of the movement trajectory that was outside the specified yellow path (presence of this red line indicated low spatial accuracy or high spatial error), and a red square indicating hand position at 650 ms (farther this red square from the target circle, lower the temporal accuracy or higher the temporal error). Based on these two performance feedback, we computed a “motor error” score which was simultaneously presented as an additional feedback at the top corner of the screen at the end of each movement. This motor error was a cumulative sum of spatial error and temporal error which were calculated using position coordinates. Data points from the red line were used to calculate the path length (in mm) that was outside the specified path (indicating spatial error). On the other hand, the red square provided hand position at 650 ms that was used to compute the distance (in mm) of the hand from the target circle (indicating temporal error). Finally, both these error values (in mm) were summed to calculate the motor error. Thus, if the participant successfully completed the movement on this skill task by reaching the target within 650 ms and within the specified path, the motor error will be close to 0. These feedback (visual cues and motor error score) were presented for 2 seconds on the screen at the end of each trial.

#### Working Memory and Control Task

After the motor skill task, participants were required to perform a secondary task for working memory engagement or a control task (based on the group type described in the next section). The working memory task used in our study was a visuo-spatial Corsi-block tapping task (Fig 1c). In this task, a blue circle (10 mm diameter) first appeared centrally on the computer screen. Upon entering the circle (with the hand-held stylus), participants heard an audio beep after 750 ms, followed by display of nine white squares. Some of these squares (3-5 out of 9) were flashed (for 750 ms each) in red color in a sequential manner. The order in which the squares changed color from white to red was random. Participants were instructed to carefully watch the flashing squares and remember the order and spatial location of the squares that changed the color. Following a second audio beep, participants were required to move their hand (displayed on screen as a yellow dot) to tap the squares that flashed red in the correct sequential order. The time to finish tapping was k+1 sec, where k was the number of squares that flashed red on each given trial (in our case 3-5 squares out of 9). After the participants finished tapping all the squares, feedback was provided on the screen as ‘Trial Successful’ if they tapped the squares in the correct sequence, otherwise ‘Trial Unsuccessful’. Subjects performed a total of 80 trials. The Corsi-block tapping task was performed with the same right arm that was used to train on the motor skill task. For subjects who performed the control task we modified this Corsi-block task such that there was no working memory requirement, and the rest of the task procedure was similar. The squares (3-5) that turned from white to red remained red and subjects were simply asked to tap all the red squares (visible on screen throughout the trial) in any order without touching the white squares. In both the control and working memory tasks, all subjects included in the analysis achieved an accuracy of 70% and above.

### Experimental Design

As mentioned in the Introduction, we conducted three experiments (with two groups of subjects in each) as part of this study. All three experiments (Exp-1, Exp-2, Exp-3) consisted of an initial training on the motor skill task with dominant right arm, followed immediately by performing a secondary task (working memory/control task) with the same trained right arm. Finally, the untrained left arm was tested for interlimb skill generalization (see Figure 1d). In Exp-1 (Long-Immediate), subjects received long training on the motor skill task (160 trials) on Day-1, then performed the secondary task (working memory, N=16 versus control group, N=16), and were immediately tested for interlimb generalization on the motor skill task (160 trials). For Exp-2 (Long-Delayed), subjects trained on the skill task (160 trials) on Day-1, then performed a secondary task (working memory, N=15 versus control group, N=15), and were tested 24 hours later for skill generalization (160 trials). Finally based on the results of Exp-1 and Exp-2, in Exp-3 (Short-Delayed), subjects received a short training on the skill task (reduced to 50 trials to prevent asymptotic phase of learning) on Day-1, then performed the secondary task (working memory, N=15 versus control group, N=15), and were tested for interlimb skill generalization (160 trials) after 24 hours on Day-2.

#### Data Analysis and Statistical Tests

Hand position data (sampled at 120 Hz) were used to assess the motor error on our skill task. Data were analyzed using Matlab R2022b and R studio. Our primary measure of interest in this study was motor skill error which was computed as a sum of spatial and temporal errors (described above and similar to our previous work of Yadav and Mutha, 2016; 2020). The composite error value (in mm) was used for assessing learning and generalization. We averaged the motor error values for five consecutive trials (based on the five directional targets presented during the task) to reduce any directional bias and obtained bin values (32 bins in Exp-1 and Exp-2 and 10 bins in Exp-3). For each bin, we also computed peak speed (cm/s) and movement preparation time (ms) taken to initiate the movement after the go audio cue (movement start time minus go audio cue time). Finally, we performed analysis of variance (ANOVA) to statistically assess learning, and interlimb generalization in all three experiments with significance level set at 0.05. Pairwise t-tests were conducted when significant main effects were observed in ANOVA and corrected with Bonferroni-holm method for multiple comparisons. We have provided omega-square for effect sizes.

### Learning and Generalization

Reduction in the skill error, as computed above, over bins indicate successful skill learning during the training session. We performed two-way repeated-measures ANOVA with bin (first, last training) as within-subject and group (working memory, control) as between-subject factors to assess learning on the skill task in all three experiments. Next, for skill generalization from right to the left arm, the first trial of the generalization session was excluded based on the findings of Wang and Sainburg (2004), who suggested that the nervous system uses the initial trial to assess relevance of prior learning in a new context. Finally, we performed ANOVA with bin (first training, first generalization) as within subject-factor and group as between-subject factor (working memory, control) in our three experiments separately.

### Strategies used for learning and generalization

Based on our past work on skill generalization (Yadav and Mutha, 2020), we were interested in understanding whether participants developed any strategy during learning and if the same strategy was transferred to the other arm when tested for generalization after engaging in the secondary task in this current study. During skill learning, reduction of motor errors on our task required participants to make fast and accurate movements. We found that participants achieved this goal by making fast movements (i.e., increasing peak speed), or preparing to move as soon as they received the go cue (decreasing movement preparation time) or both. To quantify this, we computed peak movement speed, and movement preparation time for the first and last bins of the training session and assessed if there was a significant change in these parameters. Next, we performed statistical tests to analyze if the strategy adopted during training was also transferred to the untrained arm by comparing the last bin of training and first bin of generalization session in each of the three experiments.

## RESULTS

### Engaging in working memory task after long-training on a novel motor skill does not prevent immediate or delayed generalization

In Exp-1 (Long-Immediate), participants learned to reduce the motor errors on our skill task over the course of training. Hand trajectories of one representative subject are shown in Figure 2a, where the hand position at 650 ms (denoted by square) in the first bin (first five trials) is close to the start position instead of the target circle (blue circle), and movements are curved outside the specified path. However, on the last bin of the training session, hand positions are close to the target circle, and movements are within the specified path. We found that participants in both groups (working memory, control) improved their performance (reduced motor errors) over the course of training (Figure 2b). Statistical examination revealed a significant effect of bin (first training, last training) [F[1, 30] = 305.51, p<0.001, ⍵2 = 0.20] and no group [F[1, 30] = 1.93, p = 0.17, ⍵2 = 0.0009] or bin X group interactions [F[1, 30] = 0.01, p = 0.90, ⍵2 = 0.001] suggesting similar levels of learning in both groups. Interestingly, participants in both groups exhibited interlimb generalization of learning to the untrained arm (lower motor errors) when tested immediately after engaging in the secondary task. Figure 2a shows that hand paths of the left untrained arm (first bin of generalization) are within the specified path and close to the target circle at 650 ms. We noted a significant effect of bin (first bin training, first bin generalization) [F[1, 30] = 31.09, p<0.001, ⍵2 = 0.02], but no group [F[1, 30] = 0.05, p = 0.8, ⍵2 = 0.001] or bin X group interactions [F[1, 30] = 0.64, p = 0.42, ⍵2 = 0.001] suggesting similar levels of interlimb generalization following engagement in any secondary task (Working memory or Control task, see Figure 2c).

**Figure 2:**
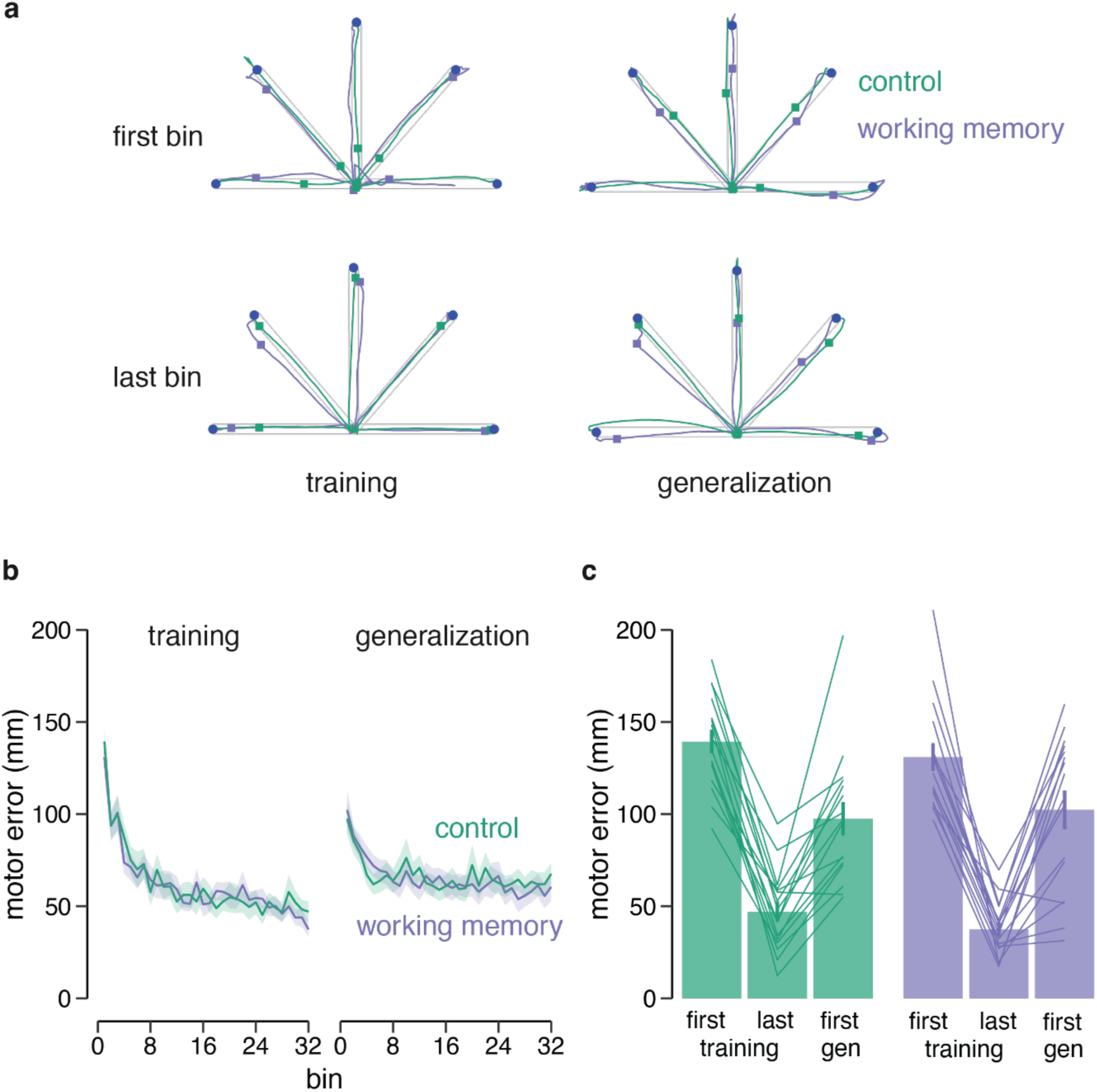
In Exp-1 (Long-Immediate), engaging in working memory or control task after long skill training does not prevent immediate interlimb generalization. a) Movement trajectories to the five targets (*blue* circle) of first and last bin of skill training and generalization sessions, respectively are shown for one participant from each group (working memory in *purple*, control in *green*). Note that as compared to the first bin of training, position of square (hand position at 650 ms) is close to the target on all other bins, indicating improved performance by the end of training and subsequent generalization of that performance to the untrained arm in participants from both groups. b) Change in mean motor error during the training and generalization sessions for the working memory (N=16) and control group (N=16). Error bands represent standard error. c) Bar plots comparing motor errors on first, last bins of training and first bin of generalization for the control (left side) and working memory (right side) groups are shown. Lines represent individual participants.

Similarly in Exp-2 (Long-Delayed), participants successfully learned the skill over the course of training session (see hand paths in Figure 3a, and group data in Figure 3b) with a significant effect of bin (first training, last training) [F[1, 28] = 323.73, p<0.001, ⍵2 = 0.22] but no group [F[1, 28] = 0.09, p = 0.75, ⍵2 = 0.0009] or bin X group interaction [F[1, 28] = 0.00005, p = 0.99, ⍵2 = 0.001]. Here too, interlimb generalization was evident when participants were tested 24 hours later-significant effect of bin [first bin training, first bin generalization] (F[1, 28] = 82.98, p<0.001, ⍵2 = 0.06), with no group [F[1, 28] = 0.24, p = 0.62, ⍵2 = 0.001] or bin X group [F[1, 28] = 0.13, p = 0.71, ⍵2 = 0.001] interactions. Thus, engaging in a visuo-spatial working memory or control task after learning a novel skill lead to similar levels of interlimb generalization to the untrained arm even at 24 hours post training (Figure 3c).

**Figure 3:**
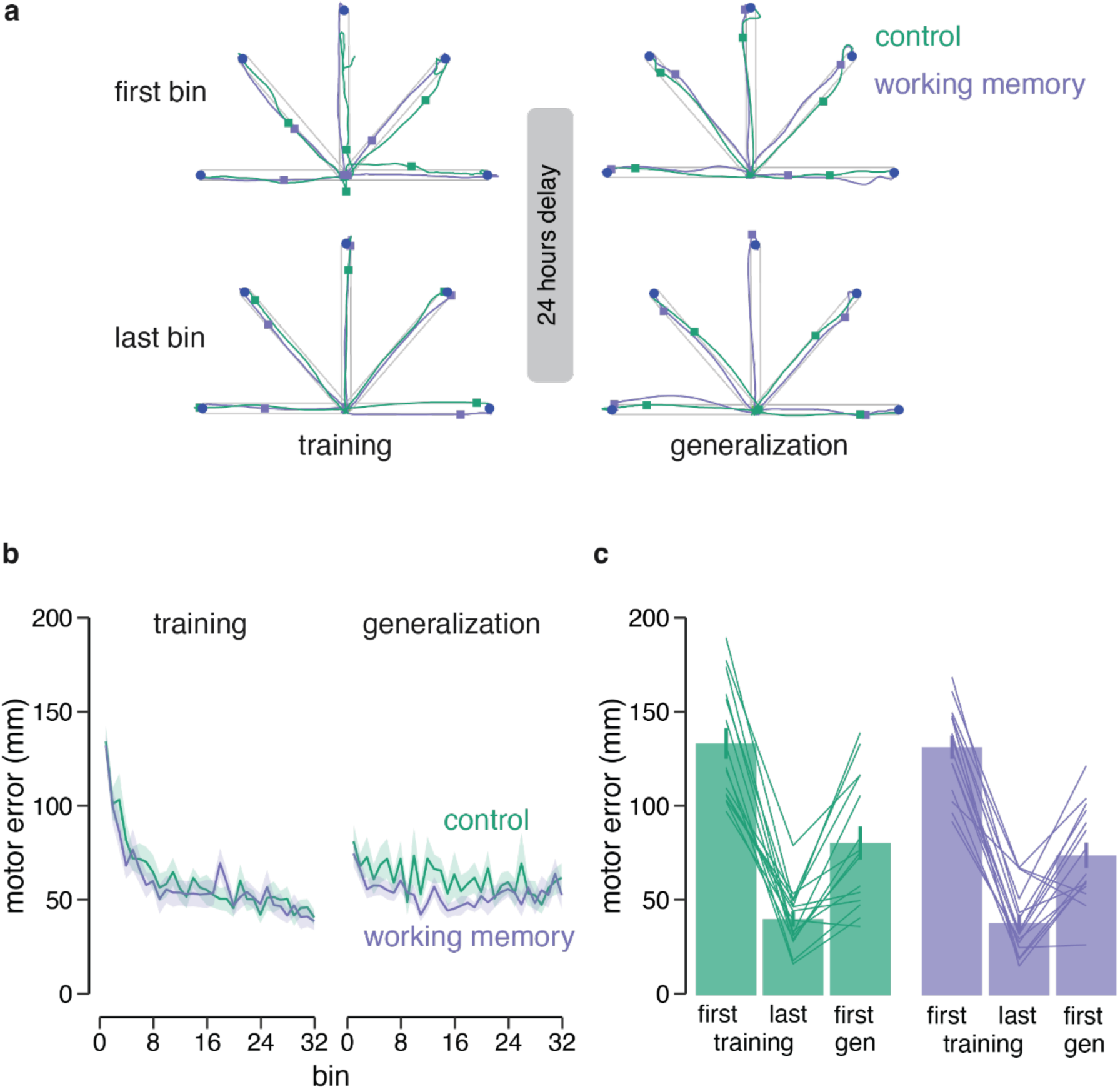
In Exp-2 (Long-Delayed), engaging in working memory or control task after long skill training does not prevent delayed interlimb generalization. a) Movement trajectories to the five targets (*blue* circle) of first and last bin of skill training and generalization sessions, respectively are shown for one participant from each group (working memory in *purple*, control in *green*). Note, like Exp-1 here too as compared to the first bin of training, position of square (hand position at 650 ms) is close to the target on all other bins, indicating learning and subsequent generalization of that performance to the untrained arm in participants of both groups. b) Change in mean motor error during training and generalization sessions for the working memory (N=15) and control group (N=15). Error bands represent standard error. c) Bar plots comparing motor errors on first, last bins of training and first bin of generalization for the control (left side) and working memory (right side) groups are shown. Lines represent individual participants.

Taken together, findings of Exp-1 and Exp-2 suggest that performing a secondary task (working memory or control task) after learning a novel skill task led to similar amounts of interlimb generalization when the untrained arm is tested immediately or 24 hours later. One plausible reason for this generalization could be the long training session with the right arm during which all participants reached the asymptotic performance around the 10th bin of the training session and repeated the learned skilled movement for the rest of the session (22 bins). We conjecture that such repetitive skill behavior may have reduced the requirement for working memory. Longer training may instead engage reinforcement mechanisms where successful actions are selected and executed without much cognitive resources making the memory hyper-stable (Shibata et al., 2017; Haith and Krakauer, 2018). To directly test this idea, we performed a third experiment (Short-Delayed) as described earlier in which we shortened the training session to 10 bins. Here again, we tested two groups (Working memory, Control) for interlimb skill generalization 24 hours after training.

### Engaging in working memory after short training on skill task impairs delayed interlimb generalization

All participants showed improvements in skill motor performance (reduced motor errors) even with the short training of 50 trials/10 bins. This reduction in the motor errors was found to be significant from first bin to last bin of the training session [F[1, 28] = 147.03, p<0.001, ⍵2 = 0.82] and similar for the two groups [F[1, 28] = 0.08, p = 0.76, ⍵2 = -0.03] with no bin X group interaction [F[1, 28] = 0.13, p=0.71, ⍵2 = -0.022]. However, unlike the previous two experiments, the working memory group exhibited impairment in interlimb generalization as compared to the Control group when tested 24 hours later (see hand paths in Figure 4a). In fact, participants in the working memory group made higher errors throughout the generalization session compared to the control group (see Figure 4b). Statistically, here we found a significant bin (first training, first generalization) X group (working memory, control) interaction (F[1, 28] = 5.10, p = 0.003, ⍵2 = 0.12). Post-hoc comparisons revealed that motor errors on the first bin of the generalization session were significantly higher in the working memory group as compared to that of the control group (*t_[28]_* = -3.5096, *p* = 0.0015), see Figure 4c.

**Figure 4:**
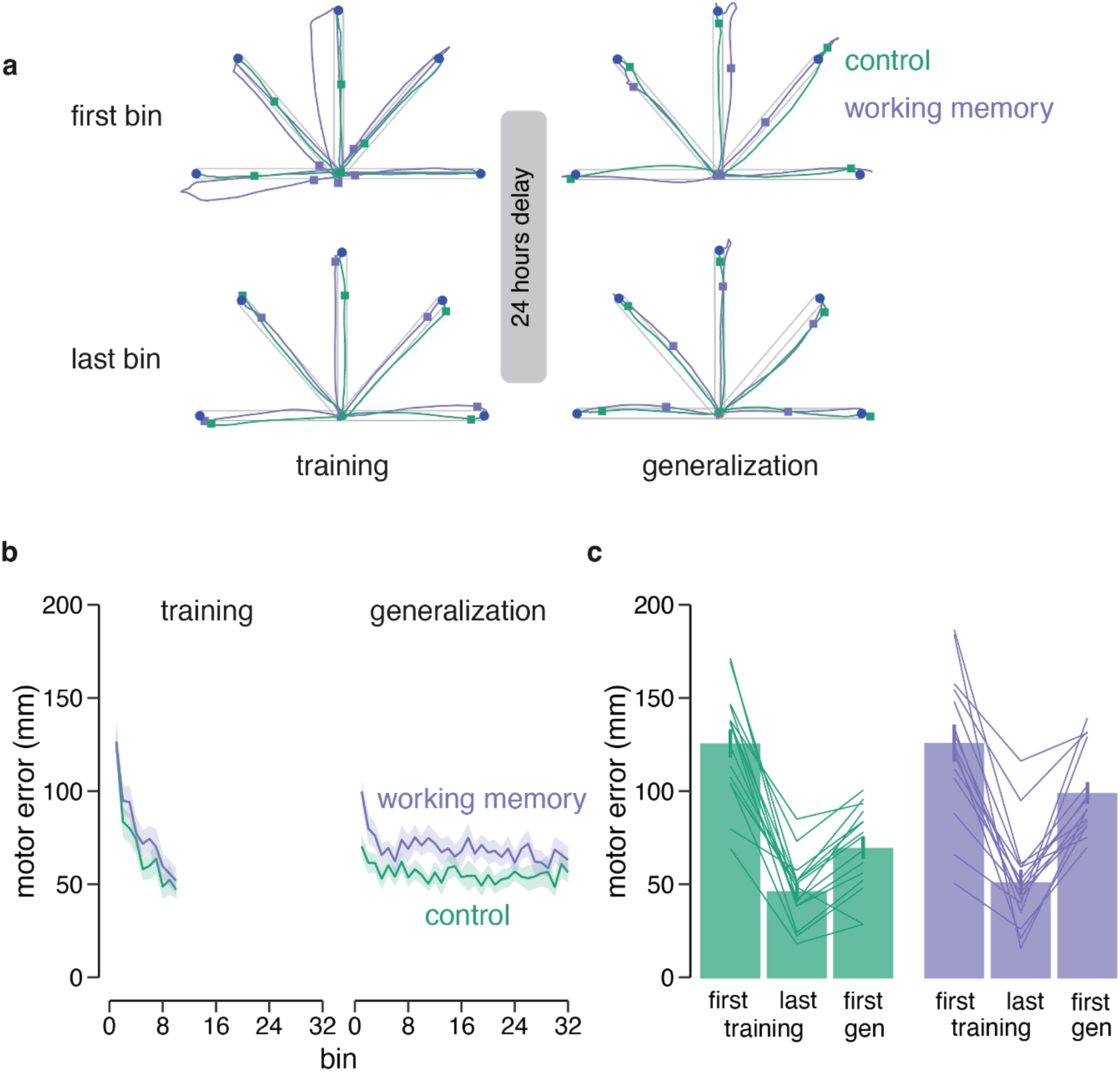
In Exp-3 (Short-Delayed), engaging in working memory but not control task after short skill training impairs interlimb generalization. a) Movement trajectories to the five targets (*blue* circle) of first and last bin of training and generalization sessions, respectively are shown for one participant from each group (working memory in *purple*, control in *green*). Note that as compared to the first bin of training, square (hand position at 650 ms) is close to the target on all other bins, indicating improved performance during training and subsequent generalization of that performance to the untrained arm in Control, but not the participant who performed working memory task following skill training. b) Change in mean motor error during the training and generalization sessions for the working memory (N=15) and control group (N=15). Error bands represent standard error. c) Bar plots comparing motor errors on first bin, last bin of training and first bin generalization for the control (left side) and working memory (right side) groups are shown. Lines represent individual participants.

Next, we wanted to understand how engaging in working memory task affected delayed interlimb generalization at 24 hours depending on the training duration. For this we pooled and directly compared the working memory groups from this experiment (Short-Delayed) and Exp-2 (Long-Delayed). We found that the amount of learning even with short training was similar in the two groups (F[1, 56] = 0.07789, p = 0.781, ⍵2 = - 0.0007). For interlimb generalization, we found that the working memory group who had received short training exhibited significantly higher motor errors on the first bin of generalization as compared to working memory group who received longer training on day-1 (*t_[28]_* = -2.8155, *p* = 0.008). This suggests that engaging in a secondary working memory task after short, but not long skill training impairs interlimb generalization when tested 24 hours later.

### Strategies adopted during skill learning and interlimb generalization

We analyzed movement speed and preparation time data to understand strategies adopted by participants to learn the motor task and whether the same strategy was transferred to the untrained arm or interfered with by the secondary task. In Exp-1 (Long-Immediate, see Figure 5a and 5b), in majority of the participants we found significant improvements over first and last bins of the training session for both parameters - increase in speed [working memory (13 out of 16): *t_[12_*_] =_ -4.3647 *p_adj_* = 0.001841; control (16 out of 16): *t_[15_*_]_ = -6.3765, *p_adj_* < 0.001] as well as reduction in preparation time [working memory (15 out of 16): *t_[14]_ =* 5.3979*, p_adj_* < 0.001; control (15 out of 16): *t_[14]_* = 4.7639, *p_adj_* < 0.001]. Participants who showed the significant change in the preparation time during training maintained their strategy when generalization to the untrained arm was tested. Their preparation time on the first bin of the generalization session was similar to the last bin of training (working memory: *t_[14]_* = 0.3131, *p_adj_* = 0.7609; control: *t_[14]_* = 0.15496, *p_adj_* = 0.8791). However, participants of both groups could not transfer the improved movement speed acquired during training to the generalization session. Speed on the first bin of the transfer was significantly lower than speed on the last bin of training (working memory: *t_[12]_* = 2.4867, *p_adj_* = 0.0286; control: *t_[15]_* = 2.7602, *p_adj_* = 0.01458). We believe this decrease in speed during generalization observed in both groups could be due to task related fatigue since all three sessions (training, secondary task and generalization) were performed in succession.

**Figure 5:**
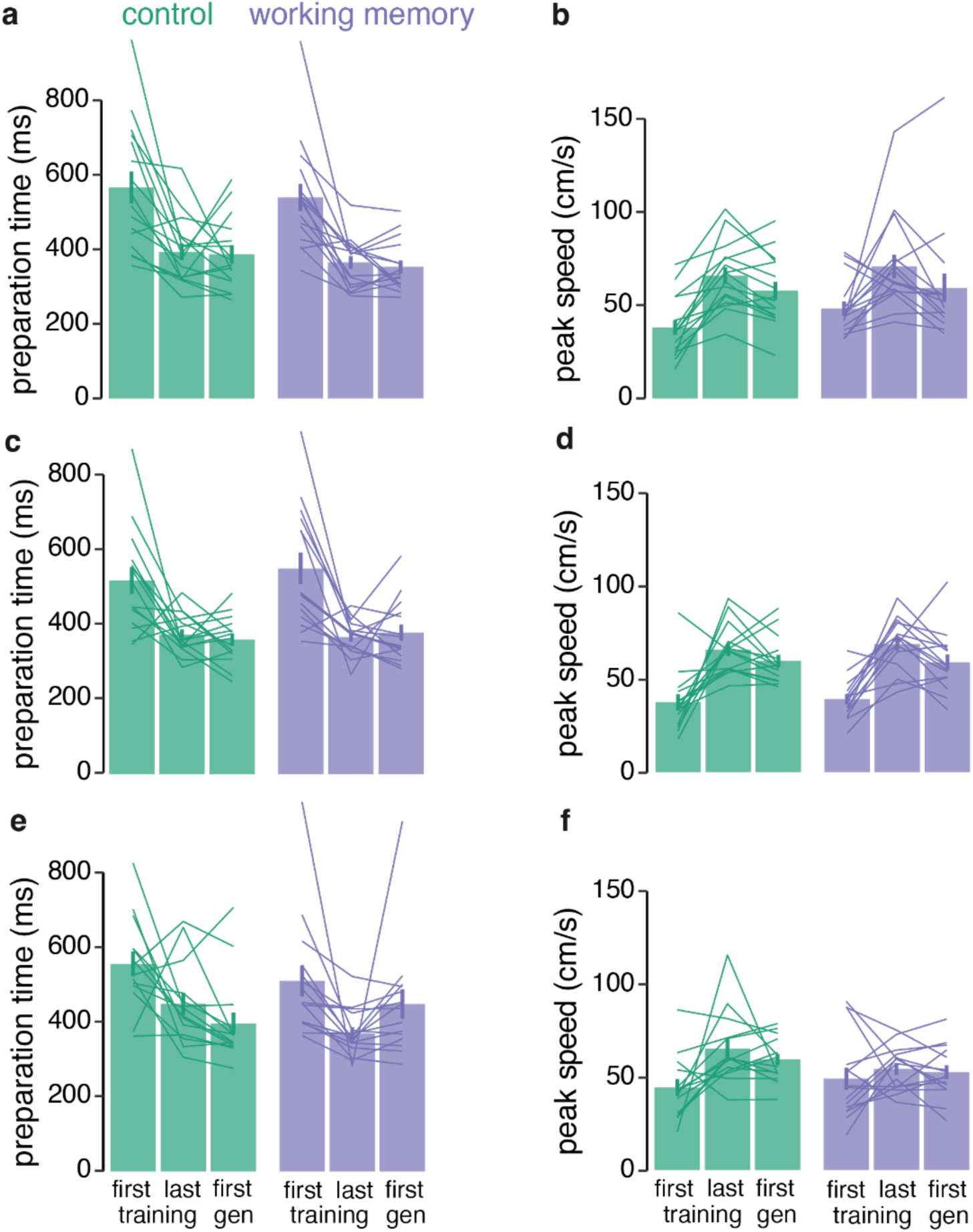
Strategy adopted during training and its subsequent generalization. For Exp-1 (Long-Immediate, *top plots*) a) mean movement preparation time and b) mean peak speed for individual participants (control indicated in *green*, working memory in *purple*) on first, last bins of training and first bin of generalization are shown. Bars represent group-averaged values. Participants reduced a) preparation time and increased b) movement speed over the first and last bin of training in both groups and transferred only improved preparation time strategy to the untrained arm when tested for generalization. In Exp-2 (Long-Delayed, *middle plots*) participants decreased c) preparation time, and d) increased speed during training with both strategies transferring to the untrained arm (see first gen) in both groups. Finally, in Exp-3 (Short-Delayed, *bottom plots*) participants in both groups decreased e) preparation time and increased f) speed during training. However, here the control group transferred both strategies to the untrained arm, but the working memory group only transferred improved speed strategy with worsening of preparation time (see first gen).

In Exp-2 (Long-Delayed, see Figure 5c and 5d), most participants also exhibited significant changes in both speed [working memory (14 out of 15): *t_[13]_* = -8.5423, *p_adj_* < 0.001; control (14 out of 15): *t_[13]_* = -5.6208, *p_adj_* < 0.001] and preparation time [working memory (14 out of 15): *t_[13]_* = 5.0196, *p_adj_* < 0.001; control (12 out of 15): *t_[11]_* = 5.2696, *p_adj_* < 0.001] during the training session. In contrast to Exp-1, both preparation time and speed strategies were transferred to the untrained arm when tested 24 hours later. Participants in both groups showed similar preparation time on the first bin of generalization and last bin of training session (working memory: *t_[13]_* = 0.55998, *p_adj_* = 0.585; control: *t_[11]_* = 0.54178, *p_adj_* = 0.5988). Similarly, speed on the first bin of the generalization session was not different from the last bin of training (working memory: *t_[13]_* = 1.9589, *p_adj_* = 0.07192; control: *t_[13]_* = 1.4585, *p_adj_* = 0.1684).

Despite reduced training, most participants in Exp-3 (Short-Delayed, see Figure 5e and 5f) also showed changes in both speed [working memory (9 out of 15): *t_[8]_* = - 5.7564, *p_adj_* < 0.001; control (11 out of 15): *t_[10]_* = -3.9344, *p_adj_* = 0.0056] as well as preparation time [working memory (14 out of 15): *t_[13]_* = 4.3515, *p_adj_* = 0.0015; control (10 out of 15): *t_[9]_* = 5.0684, *p_adj_* = 0.0013] during training. Interestingly, participants in the control group maintained their learning strategy in the generalization session when first bin of transfer was compared to last bin of training-comparable speed (t_[10]_ = 1.032, *p*_adj_ = 0.3264) with a further reduction in preparation time (*t_[9]_* = 3.6521, *p_adj_* = 0.0053). On the other hand, participants in the working memory group could not maintain the preparation time from the end of the training to the first bin of generalization session 24 hours later with worsening of preparation time (increase) in the first generalization bin as compared to the last training bin (*t_[8]_* = 2.3466, *p_adj_* = 0.0354). However, these participants maintained the same speed on the first transfer bin as in the last bin of the training (*t_[8]_* = 0.66145, *p_adj_* = 0.5269).

Overall, we found that both movement speed and preparation time underwent a significant change during training in all groups across our three experiments which helped participants learn the novel skill task. In the context of interlimb generalization, we found that the strategies that were transferred to the untrained arm depended on the experiment and group type. In Exp-1 with long training and immediate generalization test, participants in both groups transferred the movement preparation time strategy only (but not speed, possibly due to task fatigue) to the untrained arm. For Exp-2 with long training and delayed generalization test, participants in both groups transferred movement preparation time as well as speed strategies to the untrained arm. Interestingly, in Exp-3 with short training and delayed generalization test, the control group transferred both strategies of increased speed, with a further improved preparation time, whereas the working memory group transferred only speed, with worsening of preparation time. These results suggest that movement planning linked to preparation time that evolved during short training is dependent on working memory processing. On the other hand, motor control processes such as movement speed are likely not dependent and thus immune to working memory interference.

## DISCUSSION

In this study we demonstrate that engaging in a visuo-spatial working memory task as compared to control task immediately after a short skill training interferes with interlimb skill generalization tested at 24 hours. On the other hand, such generalization when tested immediately or at 24 hours is robust, and therefore not prone to any interference from working memory when the preceding skill training is long. This was surprising as the amount of learning was similar across all groups (despite long or short training) who successfully acquired the skill by adopting strategies such as increased movement speed and decreased preparation time by the end of the training session. While further exploring the specific nature of the interference caused by working memory to interlimb generalization in the short training group, we found that the strategy related to improved movement preparation time is not transferred (and in fact gets worse) to the untrained arm when it is tested at 24 hours. This suggests that engaging in a secondary cognitively demanding task engaging working memory immediately after short skill training affects skill memory stabilization in a specific manner such that the strategy related to movement preparation, but not speed, is impaired and thus not transferred to the untrained arm.

Humans have a large capacity to perform many different tasks in quick succession. However, little is known about the consequence of engaging in cognitively intensive tasks in quick succession and its impact on task memory stabilization. For instance, Cohen and Robertson (2011) show that engaging in different cognitive tasks such as skill acquisition and memorizing a list of words in quick succession can result in an underlying neural competition for memory processing. Working memory driven impairment in interlimb generalization of skill memory observed in the short skill training group is suggestive of a possible common processing bottleneck. Moreover, participants in our study were learning the skill under a variable condition which requires constant processing of the visual cues (as target location is random) before planning and executing the movement. Consequently, such learning is associated with higher cognitive load for successful performance on the task (Haith and Krakauer, 2018). In other words, variable skill learning requires trial-by-trial update of the visual cue information to aid movement execution. This working memory processing requirement is higher in the early stages of skill learning in variable task condition, such as ours, and likely more dependent on prefrontal brain areas (Kantak et., 2010; Kumar et al., 2022; Miller and Cohen, 2001).

Some prior work suggests that tasks requiring relational or associative processing elicit DLPFC activity (Smith and Jonides, 1997; Nelson et al., 2000; Robertson et al., 2001). It therefore appears that motor skill task with variable structure and the working-memory Corsi block-tapping task both share this underlying feature of relational or associative processing related to movement planning. Both these tasks, therefore, depend on DLPFC which plays an important role in initially retaining spatial information for a brief duration to allow further processing by cortical and subcortical areas to generate the desired movement (Pochon et al., 2001). In our study when participants performed the working memory task after short skill training, the stabilization/consolidation of the newly acquired motor memory was affected which resulted in impaired generalization to the untrained arm when tested the subsequent day. We propose that engaging in the working memory task likely led to an impairment in post-learning motor memory processing related to planning, as movement preparation strategy was not transferred to the untrained arm. This impairment was not observed for participants who performed the control task after short skill training, possibly because of the lack of underlying neural competition in this case. Moreover, both working memory and control groups could transfer the strategy related to speed, which we believe could be the result of motor control processes that rely on other brain areas such as the primary motor cortex, involved in processing movement kinematics in humans (Truccolo et al., 2008). This suggests that at least one component (linked to movement planning) of the newly formed motor skill memory acquired after short training, reflecting the early stage of skill learning, in our study is critically dependent on working memory processing for its further stabilization and subsequent generalization to the untrained arm.

Interestingly, long skill training resulted in significant immediate (Exp-1) as well as delayed (Exp-2) skill generalization with no interference from the subsequent working memory or control tasks. Moreover, in these experiments with long training, strategies that generalized to the untrained arm (movement preparation time transferred in Exp-1, and both movement time and speed transferred in Exp-2) did not differ for working memory or control group. We believe long training leading to the late stage of skill learning may sufficiently stabilize the newly formed skill memory with lesser reliance on working memory processing and therefore immune to interference from the secondary working memory task. This view of ours is supported by Shibata et al. (2017) who suggest that extra training after performance has reached an asymptote can make the newly acquired memory hyper-stable, thus impervious to subsequent interference from other tasks. As previously described in the introduction, skill human behavior is contingent on stable memory representations. Long training, therefore, is a simple way to make skill memory representation more stable and protect it from further interference from secondary followup tasks. Furthermore, interlimb generalization observed in all these groups (except for Exp-3 Short-Delayed working memory group) suggests that the memory representation formed during novel skill learning has effector-independent features which transfer to the untrained non-dominant arm-a phenomenon often not observed in interlimb generalization of motor adaptation (Wang and Sainburg, 2003; 2004; Kumar et al., 2018; Kumar et al., 2020). The specific nature of such newly acquired effector-independent skill memory representation(s) and how it is modulated by training or time-dependent memory stabilization processes that can influence generalization requires further detailed investigations.

Taken together, our findings show that newly learned skill can generalize to the untrained arm when tested immediately or 24 hours later following a secondary task performed after long skill training. However, such robust interlimb generalization is absent when individuals receive short skill training followed by engaging in a working memory task. Our work, therefore, highlights an underlying interaction between hitherto two distinct memory systems, i.e., skill and working memory, and the critical role of working memory in early stages of skill learning and its generalization to the untrained effector.

## ACKNOWLEDGEMENT

We would like to thank Bhoomika Sonane and Rechu Diwakar for their help with task setup.

## REFERENCES

Bao, S., & Lei, Y. (2022). Memory decay and generalization following distinct motor learning mechanisms. Journal of Neurophysiology, 128(6), 1534–1545.

Brainard, D. H., & Vision, S. (1997). The psychophysics toolbox. Spatial vision, 10(4), 433–436.

Chase, C., & Seidler, R. (2008). Degree of handedness affects intermanual transfer of skill learning. Experimental brain research, 190, 317–328.

Cohen, D. A., & Robertson, E. M. (2011). Preventing interference between different memory tasks. Nature neuroscience, 14(8), 953–955.

Criscimagna-Hemminger, S. E., Donchin, O., Gazzaniga, M. S., & Shadmehr, R. (2003). Learned dynamics of reaching movements generalize from dominant to nondominant arm. Journal of neurophysiology, 89(1), 168–176.

Fregni, F., Boggio, P. S., Nitsche, M., Bermpohl, F., Antal, A., Feredoes, E., … & Pascual-Leone, A. (2005). Anodal transcranial direct current stimulation of prefrontal cortex enhances working memory. Experimental brain research, 166, 23–30.

Haith, A. M., & Krakauer, J. W. (2018). The multiple effects of practice: skill, habit and reduced cognitive load. Current opinion in behavioral sciences, 20, 196–201.

Inui, N. (2005). Lateralization of bilateral transfer of visuomotor information in right-handers and left-handers. Journal of motor behavior, 37(4), 275–284.

Joiner, W. M., Brayanov, J. B., & Smith, M. A. (2013). The training schedule affects the stability, not the magnitude, of the interlimb transfer of learned dynamics. Journal of neurophysiology, 110(4), 984–998.

Kantak, S. S., Sullivan, K. J., Fisher, B. E., Knowlton, B. J., & Winstein, C. J. (2010). Neural substrates of motor memory consolidation depend on practice structure. Nature neuroscience, 13(8), 923–925.

Klauer, K. C., & Zhao, Z. (2004). Double dissociations in visual and spatial short-term memory. Journal of experimental psychology: General, 133(3), 355.

Krakauer, J. W., Mazzoni, P., Ghazizadeh, A., Ravindran, R., & Shadmehr, R. (2006). Generalization of motor learning depends on the history of prior action. PLoS biology, 4(10), e316.

Kumar, A., Panthi, G., Divakar, R., & Mutha, P. K. (2020). Mechanistic determinants of effector-independent motor memory encoding. Proceedings of the National Academy of Sciences, 117(29), 17338–17347.

Kumar, N., Sidarta, A., Smith, C., & Ostry, D. J. (2022). Ventrolateral prefrontal cortex contributes to human motor learning. eneuro, 9(5).

Kumar, N., Kumar, A., Sonane, B., & Mutha, P. K. (2018). Interference between competing motor memories developed through learning with different limbs. Journal of neurophysiology, 120(3), 1061–1073.

Lefumat, H. Z., Vercher, J. L., Miall, R. C., Cole, J., Buloup, F., Bringoux, L., … & Sarlegna, F. R. (2015). To transfer or not to transfer? Kinematics and laterality quotient predict interlimb transfer of motor learning. Journal of Neurophysiology, 114(5), 2764–2774.

Miller, E. K., & Cohen, J. D. (2001). An integrative theory of prefrontal cortex function. Annual review of neuroscience, 24(1), 167–202.

Mosha, N., & Robertson, E. M. (2016). Unstable memories create a high-level representation that enables learning transfer. Current Biology, 26(1), 100–105.

Nelson, C. A., Monk, C. S., Lin, J., Carver, L. J., Thomas, K. M., & Truwit, C. L. (2000). Functional neuroanatomy of spatial working memory in children. Developmental psychology, 36(1), 109.

Oldfield, R. C. (1971). The assessment and analysis of handedness: the Edinburgh inventory. Neuropsychologia, 9(1), 97–113.

Pochon, J. B., Levy, R., Poline, J. B., Crozier, S., Lehéricy, S., Pillon, B., … & Dubois, B. (2001). The role of dorsolateral prefrontal cortex in the preparation of forthcoming actions: an fMRI study. Cerebral cortex, 11(3), 260–266.

Poggio, T., & Bizzi, E. (2004). Generalization in vision and motor control. Nature, 431(7010), 768–774.

Reis, J., Schambra, H. M., Cohen, L. G., Buch, E. R., Fritsch, B., Zarahn, E., … & Krakauer, J. W. (2009). Noninvasive cortical stimulation enhances motor skill acquisition over multiple days through an effect on consolidation. Proceedings of the National Academy of Sciences, 106(5), 1590–1595.

Renault, A. G., Lefumat, H., Miall, R. C., Bringoux, L., Bourdin, C., Vercher, J. L., & Sarlegna, F. R. (2020). Individual movement features during prism adaptation correlate with after-effects and interlimb transfer. Psychological Research, 84, 866–880.

Robertson, E. M. (2009). From creation to consolidation: a novel framework for memory processing. PLoS biology, 7(1), e1000019.

Robertson, E. M. (2012). New insights in human memory interference and consolidation. Current Biology, 22(2), R66–R71.

Robertson, E. M. (2018). Memory instability as a gateway to generalization. PLoS biology, 16(3), e2004633.

Robertson, E. M., Pascual-Leone, A., & Miall, R. C. (2004). Current concepts in procedural consolidation. Nature Reviews Neuroscience, 5(7), 576–582.

Robertson, E. M., Tormos, J. M., Maeda, F., & Pascual-Leone, A. (2001). The role of the dorsolateral prefrontal cortex during sequence learning is specific for spatial information. Cerebral Cortex, 11(7), 628–635.

Sala, J. B., Rämä, P., & Courtney, S. M. (2003). Functional topography of a distributed neural system for spatial and nonspatial information maintenance in working memory. Neuropsychologia, 41(3), 341–356.

Seidler, R. D. (2004). Multiple motor learning experiences enhance motor adaptability. Journal of cognitive neuroscience, 16(1), 65–73.

Shadmehr, R., & Mussa-Ivaldi, F. A. (1994). Adaptive representation of dynamics during learning of a motor task. Journal of neuroscience, 14(5), 3208–3224.

Shibata, K., Sasaki, Y., Bang, J. W., Walsh, E. G., Machizawa, M. G., Tamaki, M., … & Watanabe, T. (2017). Overlearning hyperstabilizes a skill by rapidly making neurochemical processing inhibitory-dominant. Nature neuroscience, 20(3), 470–475.

Shmuelof, L., Krakauer, J. W., & Mazzoni, P. (2012). How is a motor skill learned? Change and invariance at the levels of task success and trajectory control. Journal of neurophysiology, 108(2), 578–594.

Smith, E. E., & Jonides, J. (1997). Working memory: A view from neuroimaging. Cognitive psychology, 33(1), 5–42.

Sun, R., Zhang, X., Slusarz, P., & Mathews, R. (2007). The interaction of implicit learning, explicit hypothesis testing learning and implicit-to-explicit knowledge extraction. Neural networks, 20(1), 34–47.

Taylor, J. A., Krakauer, J. W., & Ivry, R. B. (2014). Explicit and implicit contributions to learning in a sensorimotor adaptation task. Journal of Neuroscience, 34(8), 3023–3032.

Telgen, S., Parvin, D., & Diedrichsen, J. (2014). Mirror reversal and visual rotation are learned and consolidated via separate mechanisms: recalibrating or learning de novo?. Journal of Neuroscience, 34(41), 13768–13779.

Thoroughman, K. A., & Shadmehr, R. (2000). Learning of action through adaptive combination of motor primitives. Nature, 407(6805), 742–747.

Truccolo, W., Friehs, G. M., Donoghue, J. P., & Hochberg, L. R. (2008). Primary motor cortex tuning to intended movement kinematics in humans with tetraplegia. Journal of Neuroscience, 28(5), 1163–1178.

Wang, J., & Sainburg, R. L. (2003). Mechanisms underlying interlimb transfer of visuomotor rotations. Experimental Brain Research, 149(4), 520–526.

Wang, J., & Sainburg, R. L. (2004). Interlimb transfer of novel inertial dynamics is asymmetrical. Journal of neurophysiology, 92(1), 349–360.

Wang, J., & Sainburg, R. L. (2006). Interlimb transfer of visuomotor rotations depends on handedness. Experimental Brain Research, 175, 223–230.

Yadav, G., & Duque, J. (2023). Reflecting on what is “skill” in human motor skill learning. Frontiers in Human Neuroscience, 17, 1117889.

Yadav, G., & Mutha, P. K. (2016). Deep breathing practice facilitates retention of newly learned motor skills. Scientific reports, 6(1), 37069.

Yadav, G., & Mutha, P. K. (2020). Symmetric interlimb transfer of newly acquired skilled movements. Journal of neurophysiology, 124(5), 1364–1376.

